# Improving FnCas12a genome editing by exonuclease fusion

**DOI:** 10.1101/2020.07.07.191130

**Authors:** Yongqiang Wu, Qichen Yuan, Yufeng Zhu, Xiang Gao, Jiabao Song, Ziru Yin

## Abstract

Among current reported Cas12a orthologs, Francisella novicida Cas12a (FnCas12a) is less restricted by protospacer adjacent motif (PAM), which will help target previously inaccessible genomic sites. However, the activity of FnCas12a nuclease is relatively low or undetectable in human cells, limiting its application as desirable genome engineering tools. Here, we describe TEXT (<u>T</u>ethering <u>EX</u>onuclease <u>T</u>5 with FnCas12a), a fusion strategy that significantly increased the knockout efficiency of FnCas12a in human cells, at multiple genomic loci in three different cell lines. TEXT shows higher insertions and deletions (indels) efficiency than FnCas12a using different spacer lengths from 18nt to 23nt, in which 18nt results in highest fold increase, with up to 11 folds higher efficiency than FnCas12a. Deep sequencing shows that TEXT substantially increased the deletion frequency and deletion size at the targeted locus. TEXT enhances the activity of FnCas12a nuclease and expand its application in human cell genome engineering.

## Introduction

CRISPR (clustered regularly interspaced short palindromic repeats) Cas adaptive immune systems protect bacteria or archaea from viral invasion [1]. Leveraging CRISPR-Cas systems, a variety of genome engineering technologies have been developed, greatly accelerating the study of synthetic biology, gene therapy, diagnostics, plant engineering et al [2-6]. The development of alternative CRISPR nucleases other than commonly used Streptococcus pyogenes Cas9 (spCas9) would expand the genome engineering toolbox with new and potentially advantageous properties [7-9]. CRISPR-Cas12a system has been developed for genome editing applications with distinct features, such as recognizing T-rich PAM sequences, cleaving DNA with staggered cut distal to 5’ T-rich PAM, and the ability to process its own crRNA arrays by Cas12a [9-12]. In addition, different from Cas9 that uses crRNA/tracrRNA hybrid for DNA targeting, Cas12a only requires a single short (about 40 nucleotides) CRISPR RNA (crRNA), which is easier to be prepared [9]. Cas12a orthologs from Acidaminococcus sp. BV3L6 (As), Lachnospiraceae bacterium (Lb) have been used for genome engineering in many organisms including human cells [9-13]. However, the targeting range of AsCas12a and LbCas12a has been hindered by the strict requirement of a TTTV PAM sequence. Although several protein engineering strategies have generated new AsCas12a variants that can target non-canonical PAMs including TATV, TYCV, TTYN, VTTV, TRTV, there are still a large number of genomic sites that cannot be accessed by those variants [14, 15]. FnCas12a has been reported to use shorter and more frequently occurring PAM sequences (TTN or TTV) in vitro and in human cells [9, 16, 17], which is less restricted than AsCas12a and LbCas12a, and holds promise for targeting more currently inaccessible genomic regions. However, the nuclease activity of FnCas12a is relatively lower or unobservable in human cells [16-17]. Therefore, strategies that can improve the editing efficiency of FnCas12a will give full play to its advantages of wide targeting ability.

To date, two types of engineering strategies have been used to increase the knockout efficiency of CRISPR– Cas12a, either enhancing the nuclease activity of Cas12a by rational protein mutagenesis or improving the stability of crRNA by optimizing its secondary structures [18-21]. However, none of those approaches can fundamentally leverage the decisive step of gene knockout, the endogenous NHEJ (non-homologous end-joining) DNA repair pathway. Once the DNA double strand break (DSB) is created at target locus, NHEJ pathway will be responsible for ligating the break ends back together. Typically, NHEJ will use short homologous DNA sequences referred to as microhomologies to guide the repair, the microhomologies usually exist in the form of single-stranded overhangs at the DSB ends [25]. When the overhangs are perfectly compatible, NHEJ usually repairs the break accurately, without causing indels [22-24]. However, the imprecise NHEJ repair, which contribute to the knockout efficiency, can disrupt genes by randomly introducing unwanted indels at the ligation joint, occurring much more commonly when the overhangs are not compatible [22, 25-28]. FnCas12a has been characterized to generate staggered, perfectly compatible 5’ single strand overhangs at DSB ends [9], which is a desired substrate for error-free rather than indel involved NHEJ repair. Therefore, we reason that by resecting the 5’ single strand overhang at DSB ends, the NEHJ pathway can be biased to more frequently use the error-prone repair, thereby improving the overall knockout efficiency of FnCas12a.

Here, to improve the editing efficiency of FnCas12a in human cells, we fused T5 ssDNA exonuclease to N terminal of FnCas12a, namely TEXT, which can direct the cellular NHEJ repair more frequently into error-prone pathway, enabling higher indel efficiency than using FnCas12a alone. TEXT exhibits 2 to 3 folds higher genome editing activity than FnCas12a at all tested sites, also increases the deletion size under different spacer lengths from 18nt to 23nt, in which 18nt results in highest fold increase. Notably, when using 18nt spacer crRNA, the deletion size of TEXT is mainly 8nt or 9nt. In conclusion, we developed an improved FnCas12a system that has wide range of genome editing applications.

## Results

### Fusion of FnCas12a and 5’-3’ ssDNA exonuclease can enhance editing efficiency in HEK293T cells

Since Cas12a generates staggered DSB ends, which can be easily combined together by Watson–Crick base pairing, and thus will favor the accurate NHEJ repair pathway that does not contribute to the overall editing efficiency (Fig. 1a). To increase the gene editing efficiency of FnCas12a, we assume to bias the DSB repair pathway into imprecise NHEJ, by fusing an ssDNA exonuclease with FnCas12a that will degrade the perfectly compatible sticky ends (Fig. 1a). To test our hypothesis, we selected six ssDNA exonuclease candidates, including Artemis from human, RecJ and polA-exo from Escherichia coli, exonuclease from phage T5, phage T7 and phage λ, each of them was fused to either the N- or C-terminus of FnCas12a as candidate fusions for comparison tests of the editing efficiency (Fig. 1b). T7E1 assay showed that T5 EXO had the most significant effect on improving the indel efficiency of FnCas12a, increasing by 216% at the N-terminus of FnCas12a and 141% at the C-terminus of FnCas12a (Fig. 1c, Supplementary Figure S1). Although FnCas12a-Artemis and ecRecJ-FnCas12a showed slightly higher indel efficiency than FnCas12a, the rest of candidates did not improve the editing efficiency of FnCas12a. Considering that N-terminus fused T5 EXO-FnCas12a exhibited higher indel efficiency and smaller size than all other candidate fusions, we chose it for subsequent genome editing tests, hereafter we name it as TEXT (<u>T</u>ethering <u>EX</u>onuclease <u>T</u>5 with FnCas12a).

**Figure 1.**
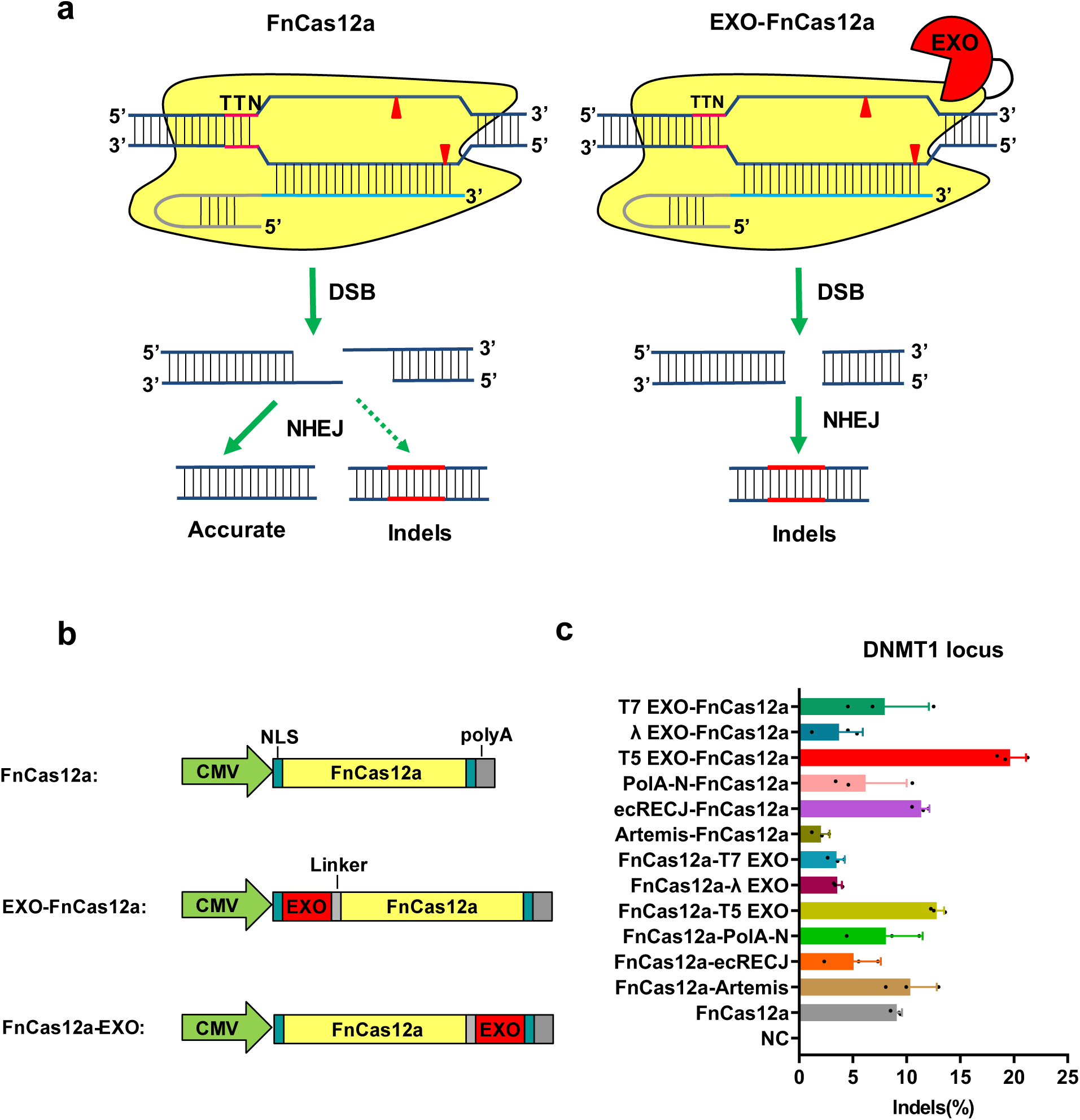
FnCas12a, EXO-FnCas12a and FnCas12a-EXO mediated gene editing in HEK293T cells. a. Illustration of the FnCas12a and EXO-FnCas12a/FnCas12a-EXO system and corresponding DNA repair pathways. FnCas12a generates fully compatible sticky ends, which are mainly repaired by precise NHEJ pathway, and thus will not contribute to indel efficiency; Exonuclease fused FnCas12a can degrade the single strand DNA overhang after cleavage, and could bias the cellular repair pathway mainly into imprecise NHEJ, which will result in the enhancement of indel efficiency. b. Schematic illustration of the FnCas12a/ EXO-FnCas12a / FnCas12a-EXO expression cassette.

### TEXT genome editing at multiple loci of different cell lines

To further profile TEXT system, we designed FnCas12a crRNAs for targeting three loci: DNMT1, CCR5 and GAPDH, in different cell lines including HEK293T cells, Hela cells and human lens epithelial B3 (HLEB3) cells. Compared to FnCas12a at DNMT1 locus, TEXT system improved gene-editing efficiency of up to 172% in Hela cells and 330% in HLEB3 cells (Fig. 2a). At CCR5 locus, TEXT showed 2 to 3 folds higher editing efficiency than FnCas12a in all three cell lines (Fig. 2b). Notably, at GAPDH locus, FnCas12a only showed around 0.3% editing efficiency in HEK293T cells, and undetectable levels of editing in both Hela and HLEB3 cells, while TEXT dramatically improved gene editing efficiency with 10-fold increase in HEK293T cells, and with significant editing efficiency in Hela and HLEB3 cells (Fig. 2c). These results indicate that TEXT system can generally increase gene editing efficiency of FnCas12a in human cells, ranging from 2 to 10 folds higher efficiency in our tests, and can significantly edit locus that previously cannot be edited by FnCas12a.

**Figure 2.**
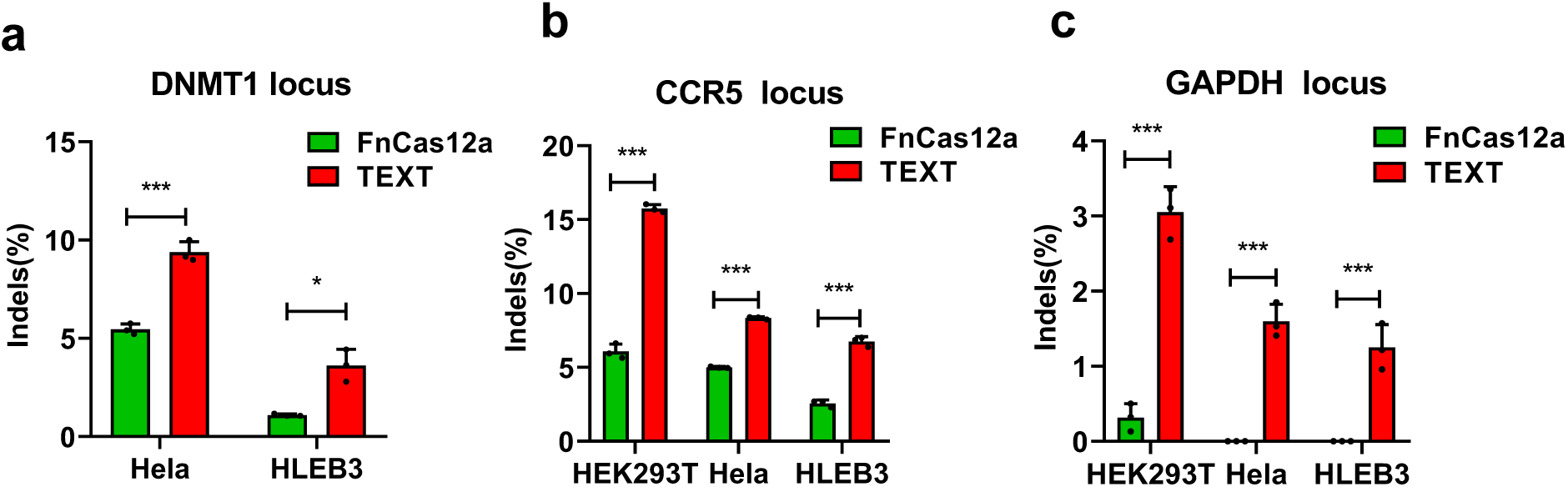
Comparison of gene editing efficiency of FnCas12a system and TEXT system in HEK293T, Hela and HLEB3 cell lines. a. Gene editing efficiency of FnCas12a system and TEXT system for the human DNMT1 gene in Hela and HLEB3 cells. b. Gene editing efficiency of FnCas12a system and TEXT system for the human CCR5 gene in HEK293T, Hela and HLEB3 cells. c. Gene editing efficiency of FnCas12a system and TEXT system for the human GAPDH gene in HEK293T, Hela and HLEB3 cells. Indel percentage at each locus was determined using the T7E1 assay and is expressed as the mean ± s.d. from three biological replicates (*P < 0.05; **P < 0.01; ***P < 0.001; two-tailed t-test). Gel images are shown in Supplementary figure S2-S4.

### Varying spacer length of crRNA to optimize editing efficiency of TEXT

The spacer region of crRNA uses Watson–Crick base-paring to locate Cas12a-crRNA complex or TEXT system at specific genomic loci. It has been reported that by optimizing the spacer length of crRNA ranging from 18nt to 23nt, the cutting efficiency of CRISPR–Cas12a can be increased, typically 21nt can achieve higher editing efficiency than other length when using FnCas12a [16]. Following a similar fashion, to investigate the optimal spacer length of crRNA for TEXT-mediated gene editing in human cells, we constructed a series of crRNAs with different spacer lengths ranging from 18nt to 23nt and tested the gene editing efficiency of TEXT at DNMT1 locus. Consistent to previous report [16], we found that crRNA with 20-21nt spacer length can enable higher gene editing efficiency when using FnCas12a (Fig. 3b), while TEXT system requires 21-23nt spacer length to achieve maximum cutting efficiency (Fig. 3b). In addition, it is also worth noting that when the spacer length of crRNA is 18nt, the editing efficiency of FnCas12a sharply decreased to 1.2%, however, TEXT system still remained more than 10% editing efficiency. Overall, these results confirmed that TEXT system is compatible with a wide range of guide length.

**Figure 3.**
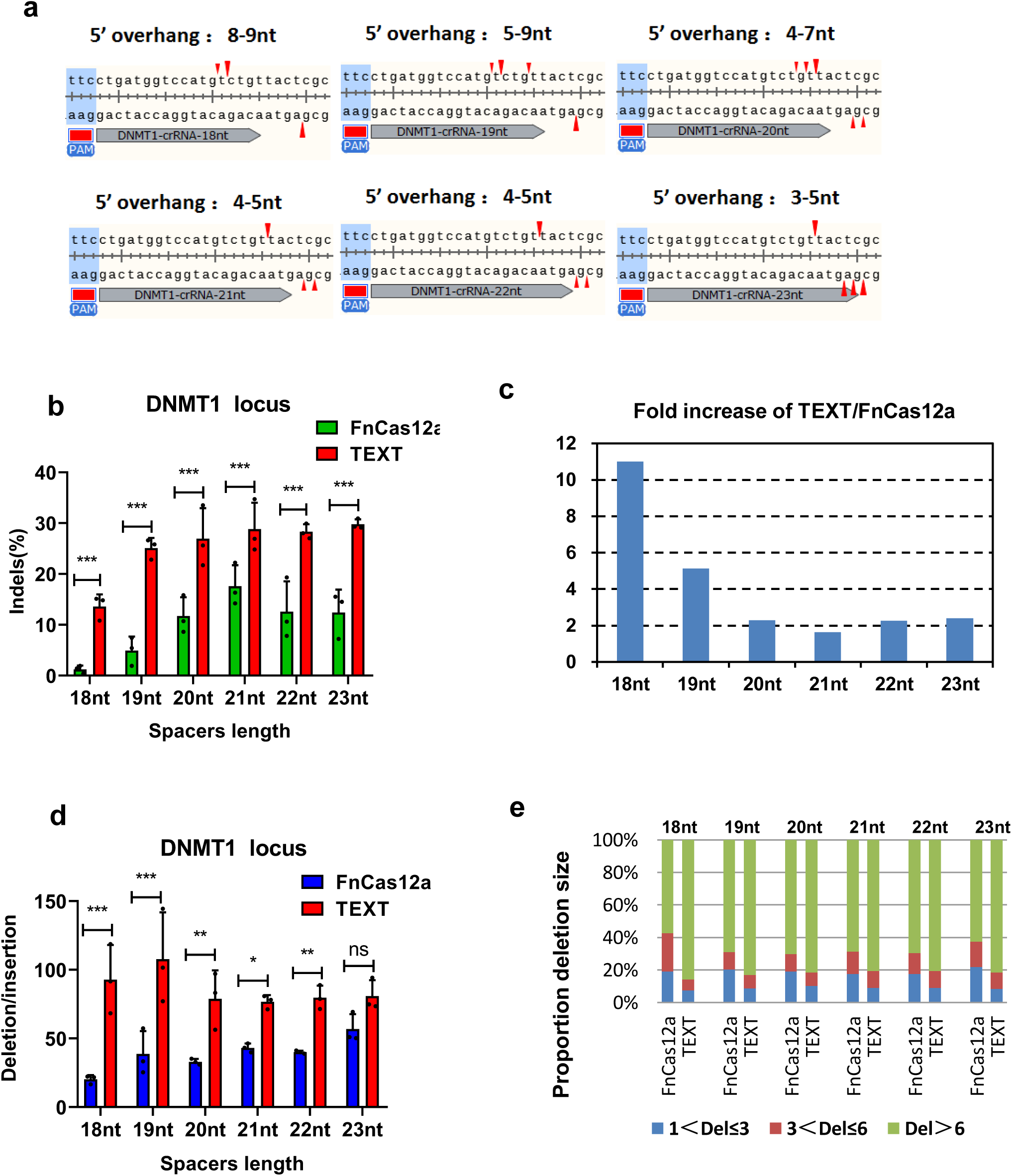
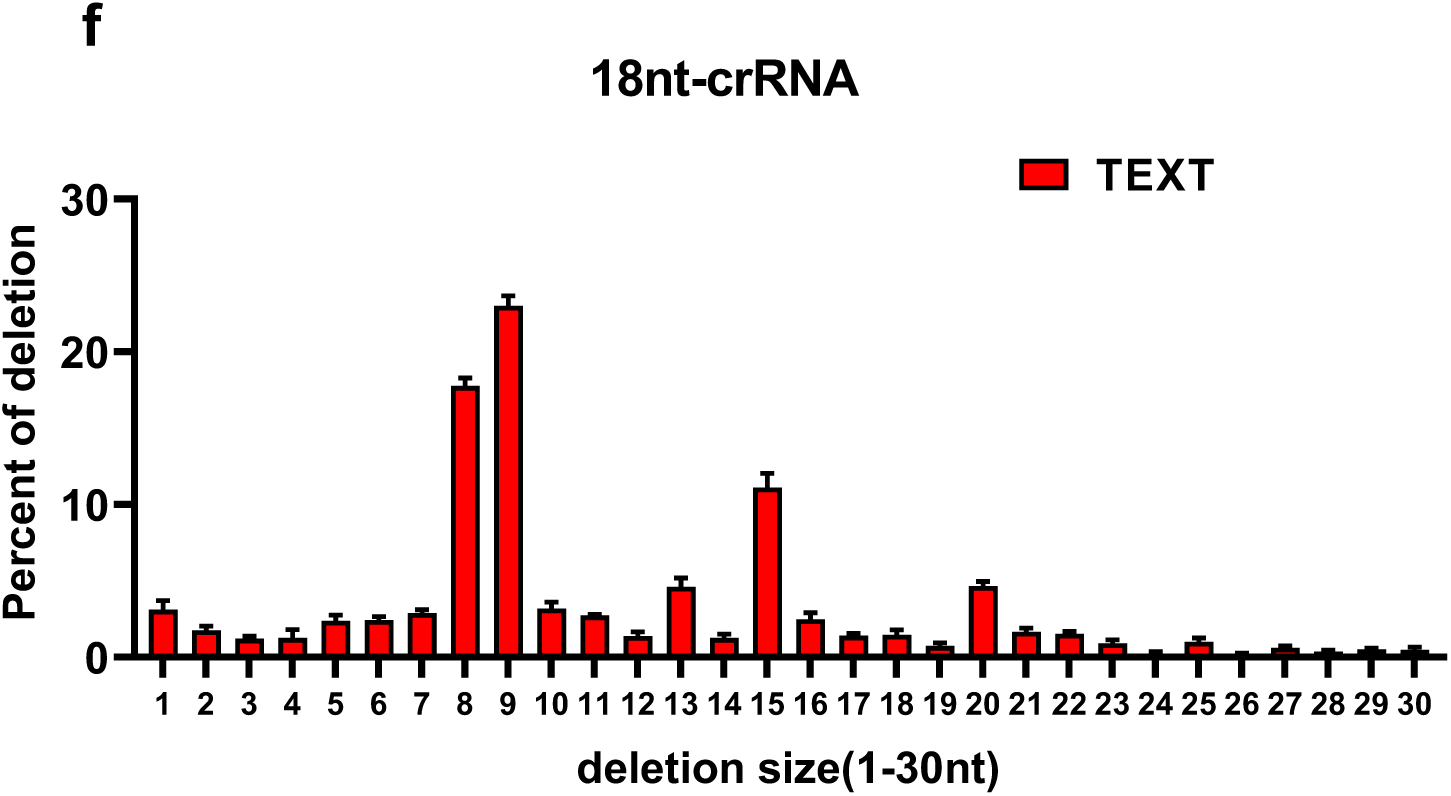
Different spacer lengths of crRNA result in distinct cleavage pattern and gene editing efficiency. a. Cleavage pattern of FnCas12a under different crRNA spacer lengths (18-23nt). With 18-nt spacer (crRNA-18 nt), FnCas12a cleavage sites were mainly after the 13th and 14th bases on the non-target strand and from the 22nd bases on the target strand. With 19-nt spacer, FnCas12a cleavage sites were mainly after the 14th,13th,17th bases on the non-target strand and from the 22nd bases on the target strand. With 20-nt spacer, FnCas12a cleavage sites were mainly after the 18th,17th,16th bases on the non-target strand and from the 18th, 19th bases on the target strand. With 21-22nt spacer, FnCas12a cleavage sites were mainly after the 18th bases on the non-target strand and from the 21st, 22nd bases on the target strand. With 23nt spacer, FnCas12a cleavage sites were mainly after the 18th bases on the non-target strand and from the 20th, 21st, 22nd bases on the target strand. b. Gene editing efficiency of FnCas12a system and TEXT system for the human DNMT1 locus with different spacer length of crRNA in HEK293T cells. Gel images are shown in Supplementary figure S5. c. Editing efficiency fold increase of TEXT system compare to FnCas12a at different spacer lengths. Ratio was determined by the formula, A/B, where A and B represent the gene editing efficiency of TEXT and FnCas12a, respectively. d. The relative ratio of deletions to insertions induced by FnCas12a or TEXT system with 18-23nt spacer lengths of crRNA. e. The relative proportions of different size deletions among total deletions that induced by FnCas12a or TEXT system with 18-23nt spacer lengths of crRNA. f. Ratio of deletions size (1-30nt) induced by TEXT system with 18nt spacer length of crRNA.

### Improving effect of TEXT under different length of crRNA

Previously, it was commonly believed that Cas12a will cut DNA at the 18th base downstream of the PAM site on the non-target strand and the 23rd base on the target strand [9]. However, recently it has been reported that FnCas12a cleavage sites are located after the 13th, 14th, 18th and 19th bases on the non-target strand and from the 21st to 24th bases on the target strand [29], which supports the scheme that FnCas12a has different cleavage pattern when using different spacer length of crRNA [29], (Fig. 3a). When the spacer length of crRNA is 18nt, which typically directs Cas12a to generate 8-9nt sticky end, the gene editing efficiency of TEXT system is 11 times higher than FnCas12a (Fig. 3a, 3c). When the spacer length is 19nt, which directs Cas12a to generate a 5-9nt sticky end, TEXT can increase the editing efficiency of FnCas12a by 5 times. (Fig. 3a, 3c). When the spacer length is 20-23bp, with 3-5nt sticky end, the gene editing efficiency of TEXT is around two-fold higher than FnCas12a (Fig. 3a, 3c).

### Deep sequencing analysis of TEXT indel patterns

Furthermore, we analyzed the indel pattern of TEXT and FnCas12a with different spacer length of crRNA by deep sequencing. (Fig. 3d-f; Supplementary Figure S6, S7). In general, TEXT induced a higher indel efficiency than FnCas12a (Fig. 3b, 3c). In addition, the ratio of deletion frequency to insertion frequency (del/in) is significantly higher in TEXT than FnCas12a (Fig. 3d). When using 18nt spacer crRNA, TEXT increased del/in of FnCas12a from 20.17 to 92.86, while only increased from 56.78 to 80.84 when using the 23nt spacer crRNA (Fig. 3d). Then we measured the lengths of deletions generated by TEXT and FnCas12a. When using FnCas12a, under different crRNA spacer length from 18nt to 23nt, the proportion of deletion size less than 3bp and less than 6bp are about 20% and 30% - 40%, respectively; while using TEXT, the deletions size that less than 3bp and less than 6bp account for 10% and 20%, respectively (Fig. 3e). It is worth noting that when the spacer length is 18nt, the deletions size that less than 6 bp is 42.6% by FnCas12a and 14.3% by TEXT (Fig. 3e). In addition, we analyzed the deletion pattern and found that the proportion of 8-9nt deletion of TEXT is up to 41.2% with 18nt spacer length crRNA, while that of FnCas12a is 9.8% (Fig. 3f). Collectively, TEXT system substantially increased the deletion frequency and deletion size at the targeted locus,and the enhancement effect will decrease when increasing the spacer length.

## Discussion

Currently, AsCas12a and LbCas12a are more widely used for genome editing than FnCas12a, although FnCas12a has shown less PAM limitations, the low targeting activity of FnCas12a limits its application as genome editing tool. We report a method that can significantly improve the editing efficiency of FnCas12a by leveraging endogenous DNA repair pathway, to favor the imprecise NHEJ repair that mainly generates indels. We also explored whether our strategy could increase the editing efficiency of LbCas12a and AsCas12a, unexpectedly, no improvement was observed on those Cas12a variants. A recent report shows that AsCas12a not only generates sticky end, but will also degrade several bases along the non-targeting ssDNA strand [30], and the structure of LbCas12a and AsCas12a are very similar, which indicates that the nuclease activities of AsCas12a and LbCas12a are high enough to approximate the saturation state of their indel efficiency, therefore it is hard to be further improved by tethering T5 exonuclease.

The crRNA of Cas12a consists of spacer sequence and direct repeat sequence that has a great impact on the editing efficiency of CRISPR/Cas12a system [9, 21]. In vitro experiments have confirmed that different spacer length of crRNA can result in different cutting patterns of FnCas12a [29]. We demonstrated that TEXT exclusively improved the gene editing efficiency of FnCas12a when using crRNAs of different spacer lengths (Fig. 3c).

In summary, TEXT significantly enhances the gene editing efficiency of FnCas12a. Our study provides insights into CRISPR–Cas12a systems and provides a new method for maximizing the genome editing efficiency of FnCas12a, enabling it as a promising human genome editing tool for broader research applications.

## Method

### Plasmid construction

PCR was performed using Q5 high-fidelity DNA polymerase (New England Biolabs) and specific primers were synthesized by Genewiz. Human codon-optimized exonuclease from phage T5, phage T7 and phage λ were synthesized by Genewiz. The Artemis gene were amplified from complementary DNA (cDNA) of HEK293T cells. Recj and PolA-exo were amplified from Escherichia coli genomic DNA. Briefly, the FnCas12a/ EXO-FnCas12a driven by CMV promoter and crRNA expression cassette driven by U6 promoter were constructed into one plasmid base on the backbone of pcDNA3.1(+) plasmid and separated by a spacer sequence. The sequences of crRNA used in this study are listed in Supplementary Table S1.

### Human cell culture and transfection

HEK293T cells and Hela cells were cultured in Dulbecco’s modified Eagle’s medium (DMEM, Gibco) supplemented with 10% fetal bovine serum (FBS, Gibco) and 1% penicillin and streptomycin (Solarbio). HLEB3 cells were cultured in DMEM supplemented with 15% FBS and 1% penicillin and streptomycin. Cells were maintained at 37 °C with 5% CO2. Cells were seeded into 96-well plates (Corning) and transfected at approximately 60% confluency. 200 ng plasmids were transfected using Lipofectamine 2000 (Life Technologies) following the manufacturer’s recommended protocol.

### Genomic DNA preparation and PCR amplification

60 hours after treatment, cells were washed with PBS genomic DNA extraction. Genomic DNA was extracted using the QuickExtract DNA Extraction Solution (Epicenter) following the manufacturer’s protocol. Concentrations of gDNA were determined on a Nanodrop 2000. Genomic regions flanking the target sites were amplified using 200 ng of purified gDNA template, Q5 high-fidelity DNA polymerase and specific primers (Supplementary Table S2) on a T100 thermal cycler (Bio-Rad)

### T7E1 cleavage assay

PCR fragments were generated using Q5 high-fidelity DNA polymerase, and then hybridized in NEBuffer 2 (50 mM NaCl, 10 mM Tris-HCl, 10 mM MgCl2, 1 mM DTT, pH 7.9) (New England Biolabs) by heating to 98 °C for 3 min, followed by a 2 °C per s ramp down to 85 °C, 1 min at 85 °C and a 0.1 °C per s ramp down to 25 °C on a T100 thermal cycler. Subsequently, the annealed samples were digested by T7 Endonuclease I (New England Biolabs) for 30 min, separated by a 2% agarose gel and quantification was based on relative band intensities. Digitalized images were analyzed to calculate indel efficiency using the Image J software. Indel percentage was determined by the formula, 100 x (1-sqrt(1-(b + c)/(a + b + c))), in which “a” represents integrated intensity of the undigested PCR product, “b” and “c” are the integrated intensities of each cleavage product, respectively.

### Deep sequencing to characterize Cas12a indels in HEK293T cells

HEK293T cells were transfected and harvested as described for assessing activity of Cas12a cleavage. The genomic-region-flanking DNMT1 targets were amplified using a pair specific primer with sample-specific barcodes added to the ends of the target amplicons (Supplementary Table S3). PCR products were run on 1.6% agarose gel and purified using Spin Column (Sangon Biotech) as per the manufacturer’s recommended protocol. Equal amounts of the PCR products were pooled, and samples were sequenced commercially (GENEWIZ, Suzhou, China) by paired-end read sequencing using the Illumina HiSeq X Ten platform.

### Statistics

Statistical significance was calculated using MannWhitney tests using GraphPad Prism version 8.0. The error bars in all figures show standard error of the mean (n = 3). P values are reported using GraphPad style: not significant (ns), P > 0.05; *, P < 0.05; **, P < 0.01; ***, P < 0.001.

## Abbreviations

Cas: CRISPR-associated protein
crRNA: CRISPR RNA
ssDNA: Single-stranded DNA
NLS: Nuclear localization signal
nt: Nucleotide

## Author contributions

Y. W. conceived the project and designed experiments. Y.W., Y.Z., J.S., X.G., Z.Y., performed the experiments. Y.W., Q.Y., analyzed data. Y.W., Q.Y., wrote the manuscript with help from all authors.

## Availability of data and materials

Y. W. is responsible for distributing plasmids and materials of this work upon reasonable request. Data sets from high-throughput sequencing experiments have been deposited with the National Center for Biotechnology Information Sequence Read Archive under BioProject ID:PRJNA642811

## References

1. Carter J, Wiedenheft B. CRISPR-RNA-guided adaptive immune systems. Cell. 2015; 163(1):260–260.

2. Doudna JA. The promise and challenge of therapeutic genome editing. Nature. 2020; 578(7794):229–236.

3. Yin, K., Gao, C. & Qiu, J.-L. Progress and prospects in plant genome editing. Nat. Plants 2017;3:17107.

4. Yin, H., Kauffman, K. J. & Anderson, D. G. Delivery technologies for genome editing. Nat. Rev. Drug Discov. 2017; 16, 387–399.

5. Wang H, La Russa M, Qi LS. CRISPR/Cas9 in Genome Editing and Beyond. Annu Rev Biochem. 2016; 85:227–64.

6. Jacinto FV, Link W, Ferreira BI. CRISPR/Cas9-mediated genome editing: From basic research to translational medicine. J Cell Mol Med. 2020; Epub ahead of print..

7. Cong, L., Ran, F.A., Cox, D., Lin, S., Barretto, R., Habib, N., Hsu, P.D., Wu, X., Jiang, W. Marraffini, L.A., and Zhang, F. Multiplex genome engineering using CRISPR/Cas systems. Science 2013; 339, 819–823.

8. Mali P, Yang L, Esvelt KM, Aach J, Guell M, DiCarlo JE, Norville JE, Church GM. RNA-guided human genome engineering via Cas9. Science. 2013; 339(6121):823–6.

9. Zetsche B, Gootenberg JS, Abudayyeh OO, Slaymaker IM, Makarova KS, Essletzbichler P, et al. Cpf1 is a single RNA guided endonuclease of a class 2 CRISPR Cas system. Cell. 2015;163(3):759–71.

10. Kim D, Kim J, Hur JK, Been KW, Yoon SH, Kim JS. Genome-wide analysis reveals specificities of Cpf1 endonucleases in human cells. Nat Biotechnol. 2016; 34(8):863–8.

11. Kleinstiver, B.P., Tsai, S.Q., Prew, M.S., Nguyen, N.T., Welch, M.M., Lopez, J.M., McCaw, Z.R., Aryee, M.J. and Joung, J.K. Genome-wide specifcities of CRISPR–Cas Cpf1 nucleases in human cells. Nat. Biotechnol. 2016; 34, 869–874.

12. Fonfara I, Richter H, Bratovič M, Le Rhun A, Charpentier E. The CRISPR-associated DNA-cleaving enzyme Cpf1 also processes precursor CRISPR RNA. Nature. 2016; 532(7600):517–21.

13. Zetsche B, Heidenreich M, Mohanraju P, Fedorova I, Kneppers J, DeGennaro EM, Winblad N Choudhury SR, Abudayyeh OO, Gootenberg JS1, Wu WY. et al. Multiplex gene editing by CRISPR-Cpf1 using a single crRNA array. Nat Biotechnol. 2017;35(1):31–34.

14. Linyi Gao, David B T Cox, Winston X Yan, John C Manteiga, Martin W Schneider, Takashi Yamano, Hiroshi Nishimasu, Osamu Nureki, Nicola Crosetto, Feng Zhang. Engineered Cpf1 variants with altered PAM specifcities. Nat. Biotechnol. 2017; 35, 789–792.

15. Kleinstiver BP, Sousa AA, Walton RT, Tak YE, Hsu JY, Clement K, Welch MM, Horng JE, Malagon-Lopez J, Scarfò I, et al. Engineered CRISPR-Cas12a variants with increased activities and improved targeting ranges for gene, epigenetic and base editing. Nat Biotechnol. 2019;37(3):276–282.

16. Tu, M., Lin, L., Cheng, Y., He, X., Sun, H., Xie, H., Fu, J., Liu, C., Li, J., Chen, D., et al. A ‘new lease of life’: FnCpf1 possesses DNA cleavage activity for genome editing in human cells. Nucleic Acids Res. 2017;45, 11295–11304.

17. Eszter Tóth, Bernadett C Czene, Péter I Kulcsár, Sarah L Krausz, András Tálas, Antal Nyeste, Éva Varga, Krisztina Huszár, Nóra Weinhardt, Zoltán Ligeti. et al. Mb - And FnCpf1 Nucleases Are Active in Mammalian Cells: Activities and PAM Preferences of Four Wild-Type Cpf1 Nucleases and of Their Altered PAM Specificity Variants. Nucleic Acids Res. 2018;46(19):10272–10285.

18. Bin Moon S, Lee JM, Kang JG, Lee NE, Ha DI, Kim DY, Kim SH1, Yoo K, Kim D, Ko JH, Kim YS. Highly efficient genome editing by CRISPR-Cpf1 using CRISPR RNA with a uridinylate-rich 3’-overhang. Nat Commun. 2018;9(1):3651.

19. Park HM, Liu H, Wu J, Chong A, Mackley V, Fellmann C, Rao A, Jiang F, Chu H, Murthy N, Lee K. Extension of the crRNA enhances Cpf1 gene editing in vitro and in vivo. Nat Commun. 2018;9(1):3313.

20. Li B, Zhao W, Luo X, Zhang X, Li C, Zeng C, Dong Y. Engineering CRISPR-Cpf1 crRNAs and mRNAs to maximize genome editing efficiency. Nat Biomed Eng. 2017;1(5):0066.

21. Li Lin 1, Xiubin He, Tianyuan Zhao, Lingkai Gu, Yeqing Liu, Xiaoyu Liu, Hongyan Liu, Fayu Yang, Mengjun Tu, Lianchao Tang. et al. Engineering the Direct Repeat Sequence of crRNA for Optimization of FnCpf1-Mediated Genome Editing in Human Cells. Mol Ther. 2018;26(11):2650–2657.

22. J K Moore, J E Haber. Cell Cycle and Genetic Requirements of Two Pathways of Nonhomologous End-Joining Repair of Double-Strand Breaks in Saccharomyces Cerevisiae. 1996 May;16(5):2164–73.

23. Joe Budman, Gilbert Chu. Processing of DNA for Nonhomologous End-Joining by Cell-Free Extract. EMBO J. 2005;24(4):849–60.

24. Jiafeng Gu, Haihui Lu, Brigette Tippin, Noriko Shimazaki, Myron F Goodman, Michael R Lieber. XRCC4:DNA Ligase IV Can Ligate Incompatible DNA Ends and Can Ligate Across Gaps. EMBO J. 2007;26(4):1010–23.

25. Michael R Lieber. The Mechanism of Double-Strand DNA Break Repair by the Nonhomologous DNA End-Joining Pathway. Annu Rev Biochem. 2010;79:181–211.

26. Mireille Bétermier, Pascale Bertrand, Bernard S Lopez. Is Non-Homologous End-Joining Really an Inherently Error-Prone Process? PLoS Genet. 2014;10(1):e1004086.

27. Markus Löbrich, Penny Jeggo. A Process of Resection-Dependent Nonhomologous End Joining Involving the Goddess Artemis. Trends Biochem Sci. 2017;42(9):690–701.

28. Chang HH, Lieber MR. Structure-Specific nuclease activities of Artemis and the Artemis: DNA-PKcs complex. Nucleic Acids Res. 2016;44(11):4991–7.

29. Chao Lei, Shi-Yuan Li, Jia-Kun Liu, Xuan Zheng, Guo-Ping Zhao, Jin Wang. The CCTL (Cpf1-assisted Cutting and Taq DNA Ligase-Assisted Ligation) Method for Efficient Editing of Large DNA Constructs in Vitro. Nucleic Acids Res. 2017;45(9):e74.

30. Joshua C Cofsky, Deepti Karandur, Carolyn J Huang, Isaac P Witte, John Kuriyan, Jennifer A Doudna. CRISPR-Cas12a Exploits R-loop Asymmetry to Form Double-Strand Breaks. Elife. 2020;9:e55143.

