## Supplementary material for "Improving FnCas12a genome editing by exonuclease fusion": bio sup V2.docx

**Supplementary Data**

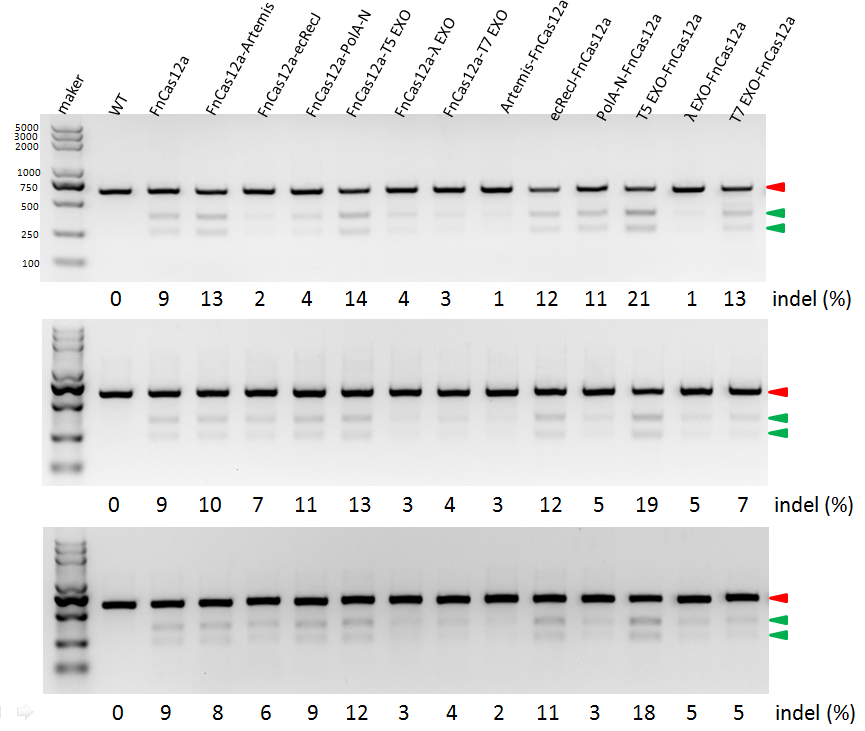

**Figure S1.** FnCas12a, EXO- FnCas12a and FnCas12a-EXO mediated gene editing efficiency in HEK293T cells. Indel percentage at each locus was determined using the T7E1 assay by quantification of the uncut (red arrow) and cut DNA bands (green arrow).

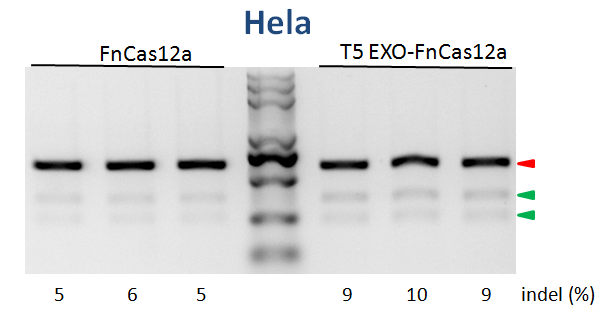

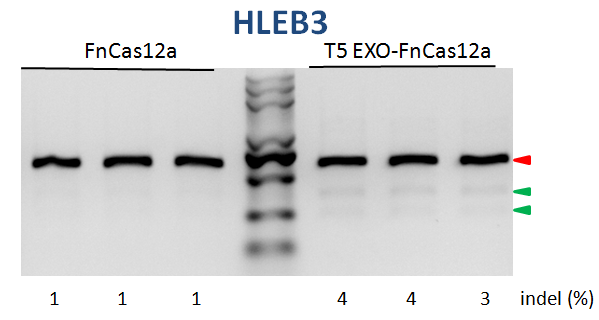

**Figure S2. FnCas12a system and TEXT system mediated gene editing efficiency for the human DNMT1 gene in Hela and HLEB3 cells.** Indel percentage at each locus was determined using the T7E1 assay by quantification of the uncut (red arrow) and cut DNA bands (green arrow).

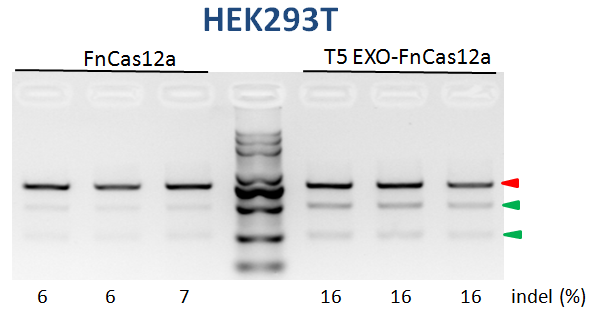

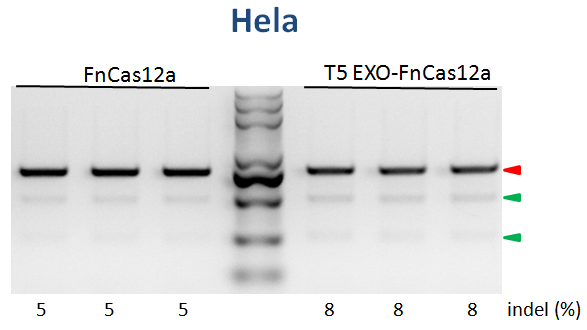

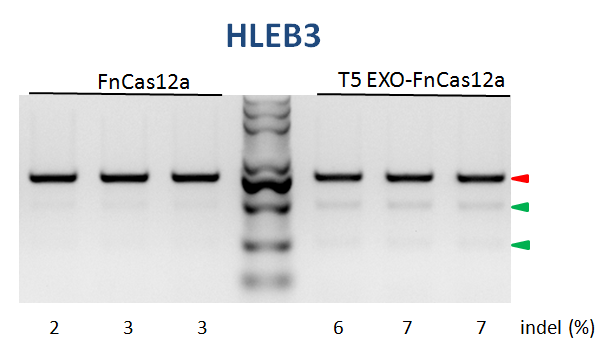

**Figure S3. Gene editing efficiency of FnCas12a system and TEXT system for the human CCR5 gene in HEK293T, Hela and HLEB3 cells.** Indel percentage at each locus was determined using the T7E1 assay by quantification of the uncut (red arrow) and cut DNA bands (green arrow).

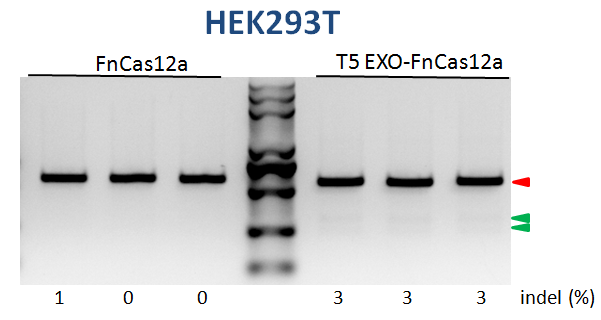

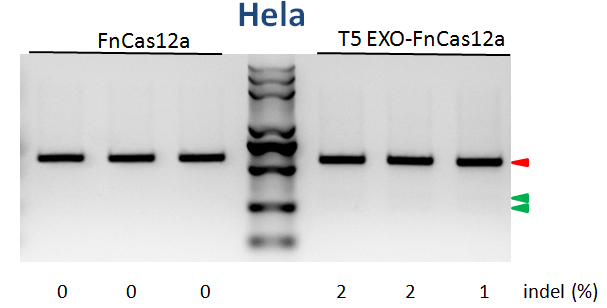

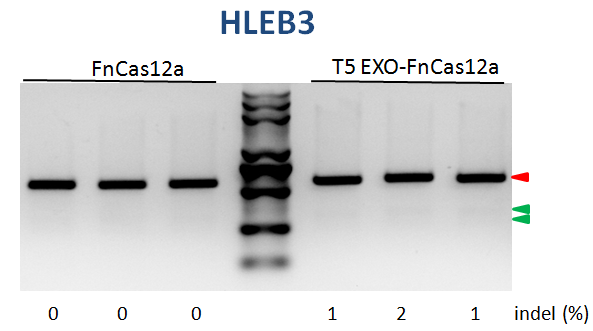

**Figure S4. Gene editing efficiency of FnCas12a system and TEXT system for the human GAPDH gene in HEK293T, Hela and HLEB3 cells.** Indel percentage at each locus was determined using the T7E1 assay by quantification of the uncut (red arrow) and cut DNA bands (green arrow).

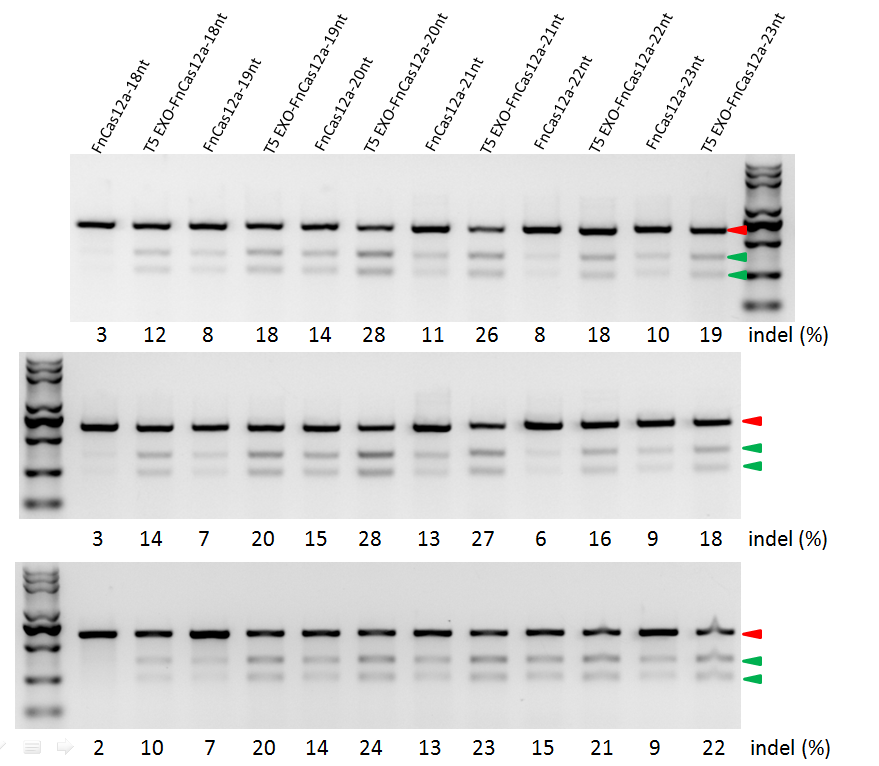

**Figure S5. Maximizing efficiency by regulate the spacer length of crRNA at DNMT1 locus.** Indel percentage at each locus was determined using the T7E1 assay by quantification of the uncut (red arrow) and cut DNA bands (green arrow).

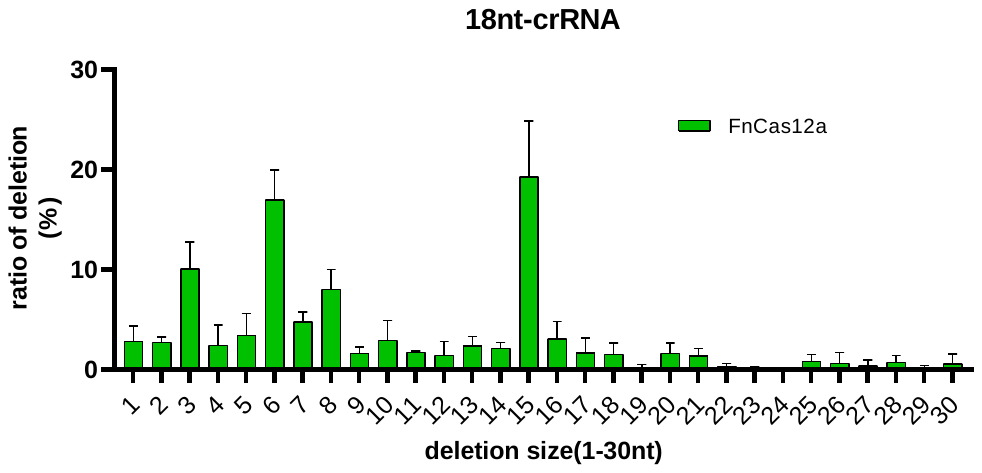

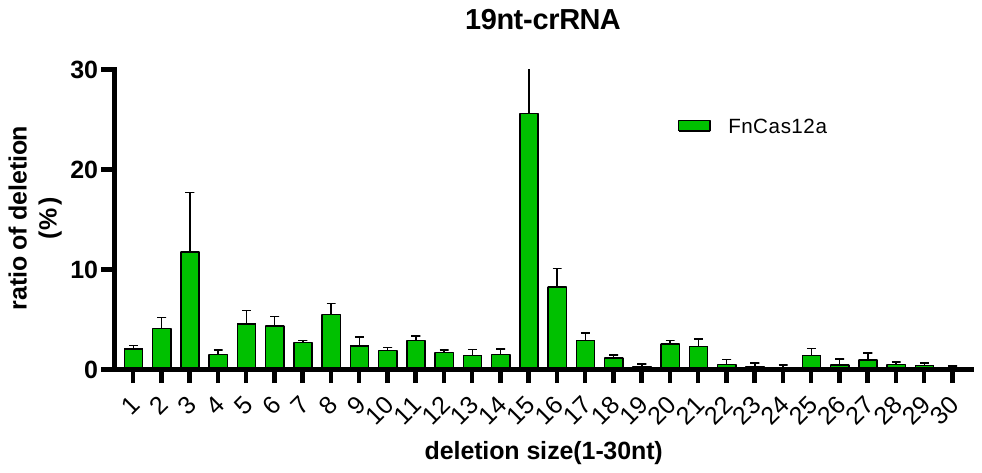

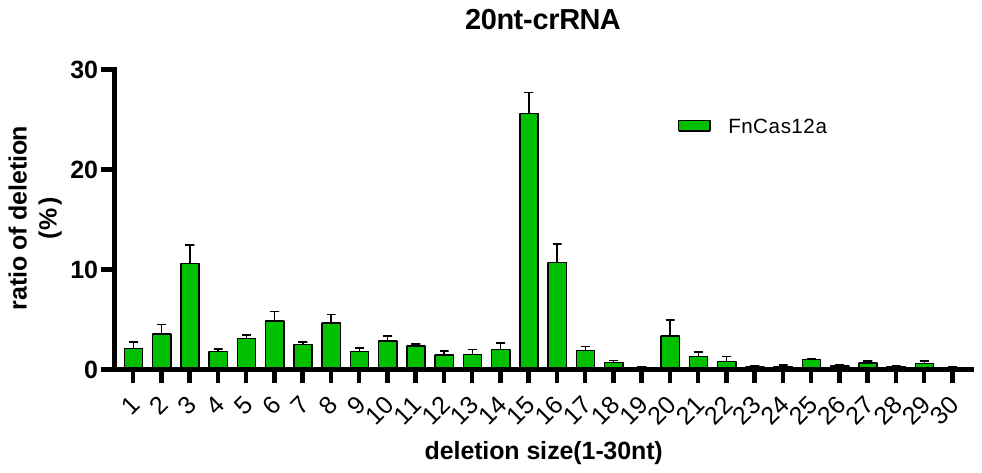

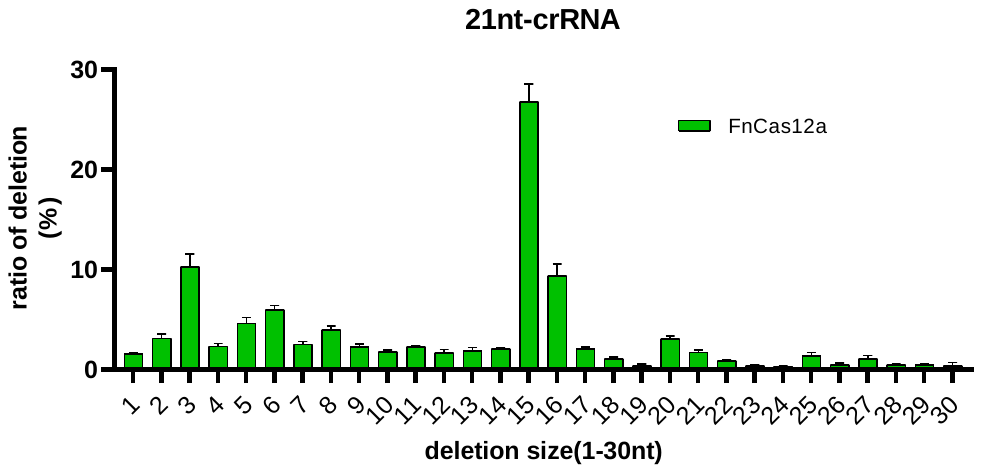

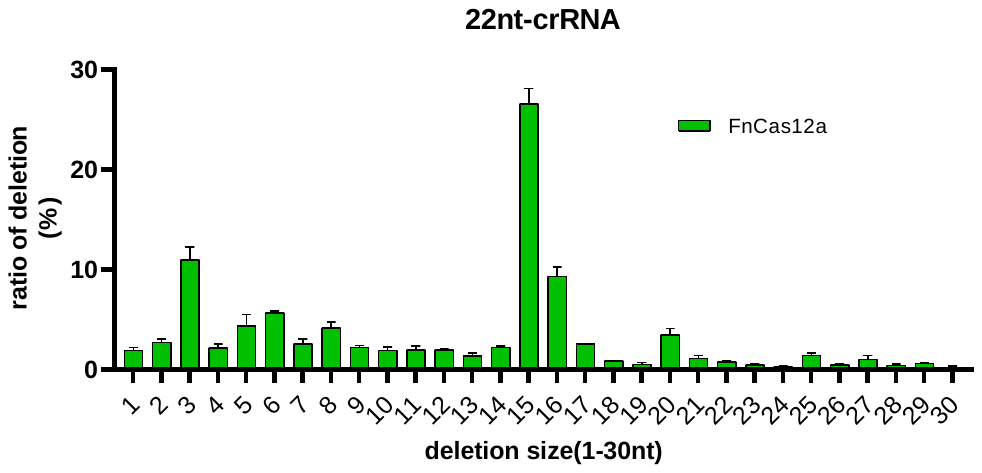

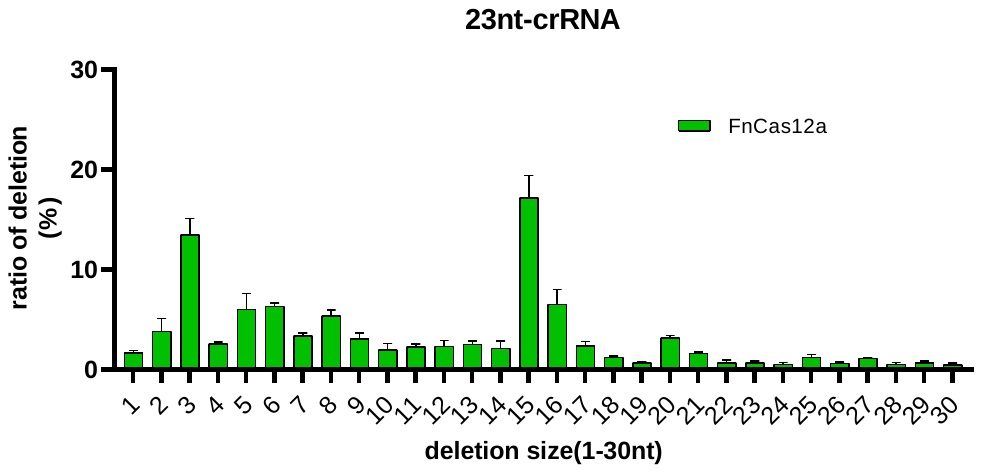
**Figure S6. Ratio of deletions size (1-30nt) induced by FnCas12a System with 18-23nt spacer length of crRNA.**

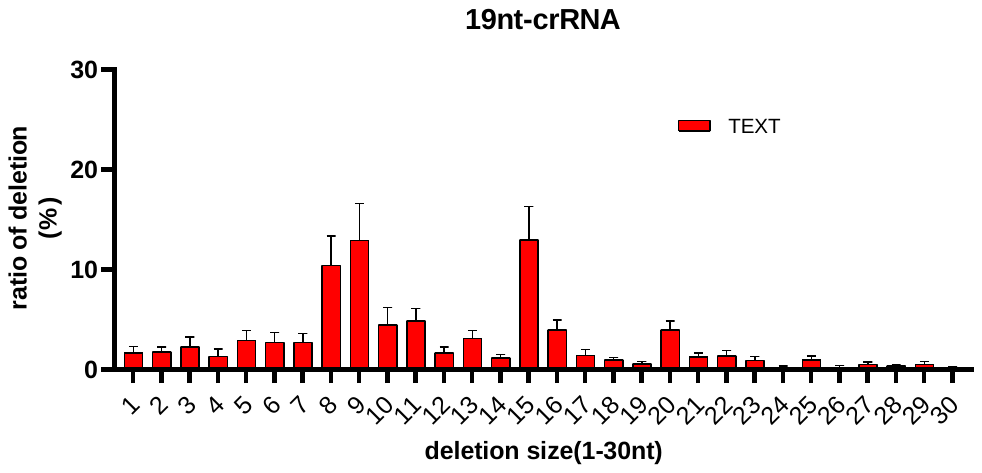

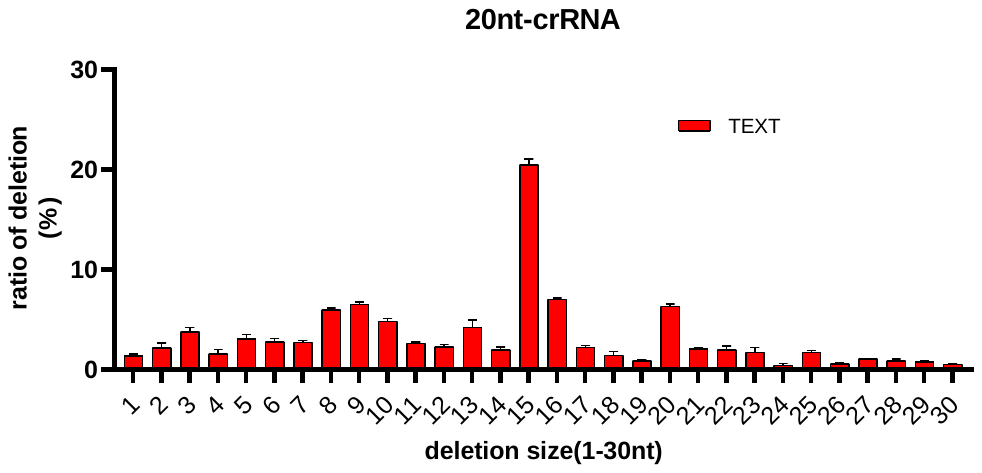

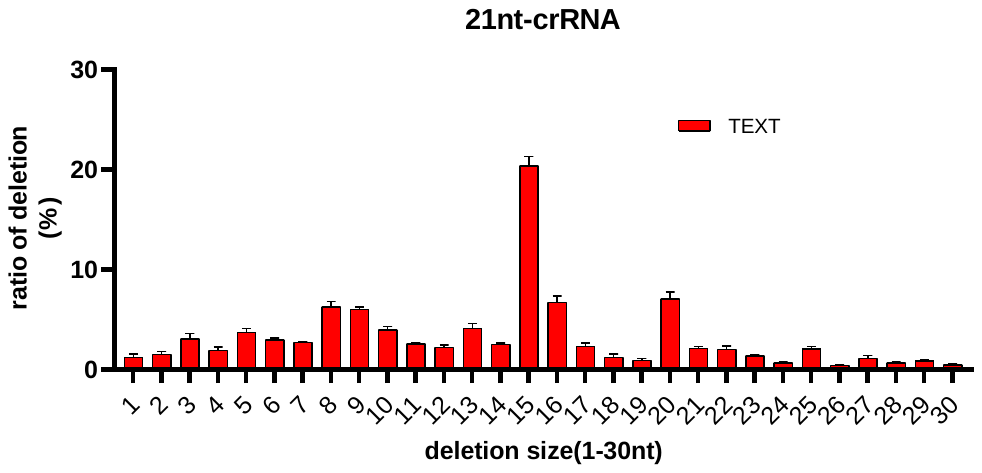

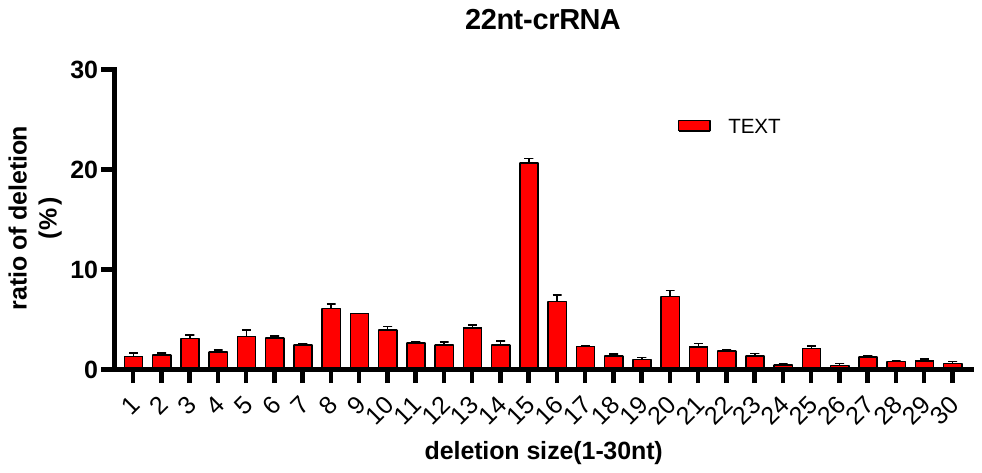

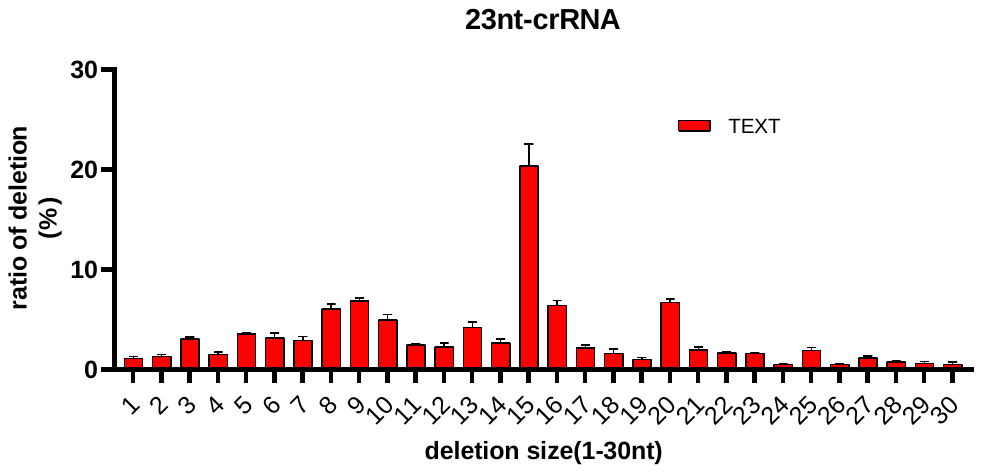

**Figure S7. Ratio of deletions size (1-30nt) induced by TEXT System with 19-23nt spacer length of crRNA.**

**Table S1. Different crRNA sequences with FnCas12a**

| **Description** | **sequences** |
| --- | --- |
| **FnCas12a crRNA at DNMT1-18nt** | ggaCGAATTTCTACTGTTGTAGATctgatggtccatgtctgtCTTTTTTC |
|  | TCGAGAAAAAAGacagacatggaccatcagATCTACAACAGTAGAAATTCGtccgc |
| **FnCas12a crRNA at DNMT1-19nt** | ggaCGAATTTCTACTGTTGTAGATctgatggtccatgtctgttTTTTTTC |
|  | TCGAGAAAAAAaacagacatggaccatcagATCTACAACAGTAGAAATTCGtccgc |
| **FnCas12a crRNA at DNMT1-20nt** | *ggaCGAATTTCTACTGTTGTAGATctgatggtccatgtctgttaTTTTTTC* |
|  | TCGAGAAAAAAtaacagacatggaccatcagATCTACAACAGTAGAAATTCGtccgc |
| **FnCas12a crRNA at DNMT1-21nt** | ggaCGAATTTCTACTGTTGTAGATctgatggtccatgtctgtTACATTTTTTC |
|  | TCGAGAAAAAATGTAacagacatggaccatcagATCTACAACAGTAGAAATTCGtccgc |
| **FnCas12a crRNA at DNMT1-22nt** | ggaCGAATTTCTACTGTTGTAGATctgatggtccatgtctgtTACTTTTTTTC |
|  | TCGAGAAAAAAAGTAacagacatggaccatcagATCTACAACAGTAGAAATTCGtccgc |
| **FnCas12a crRNA at DNMT1-23nt** | ggaCGAATTTCTACTGTTGTAGATctgatggtccatgtctgtTACTCTTTTTTC |
|  | TCGAGAAAAAAGAGTAacagacatggaccatcagATCTACAACAGTAGAAATTCGtccgc |
| **FnCas12a crRNA at CCR5** | ggaCGAATTTCTACTGTTGTAGATctgctccccagtggatcgggtgtTTTTTTC |
|  | TCGAGAAAAAAtaacagacatggaccatcagATCTACAACAGTAGAAATTCGtccgc |
| **FnCas12a crRNA at GAPDH** | ggaCGAATTTCTACTGTTGTAGATctccttggaggccatgtgggTTTTTTC |
|  | TCGAGAAAAAAcccacatggcctccaaggagATCTACAACAGTAGAAATTCGtccgc |

**Table S2. Primers used for T7E1 assay.**

| Locus | Forward Primers (5’-3’) | Reverse Primers (5’-3’) |
| --- | --- | --- |
| DNMT1-1 | CTGGGACTCAGGCGGGTCAC | CCTCACACAACAGCTTCATGTCAGC |
| CCR5 | CCTTCTCCTGAACACCTTCC | CCATAGCAAGACAAAGACCTG |
| GAPDH | GTGGTCTCCTCTGACTTCAAC | CCAGCAAGAATGTCTCACC |

**Table S3 A list of primers used for the 1st round of PCR for deep sequencing.** Specific barcode sequences are in red.

| **ID.** | **Sample** | **PCR primers** | **Index** |
| --- | --- | --- | --- |
| A | FnCas12a-18nt  Repeat1 | F1: ATCGATTCCCTCACTCCTGCTCGGTGAA  R1: AGTAGACCAAGTCACTCTGGGGAACACGCC | CTGAAGCT+CACAGGAT |
| A | FnCas12a-18nt  Repeat2 | F2: CGATGTTCCCTCACTCCTGCTCGGTGAA  R2: TTATCTATAAGTCACTCTGGGGAACACGCC | CTGAAGCT+CACAGGAT |
| A | FnCas12a-18nt  Repeat3 | F3: TTAGGCTCCCTCACTCCTGCTCGGTGAA  R3: CATGACATAAGTCACTCTGGGGAACACGCC | CTGAAGCT+CACAGGAT |
| B | T5 EXO-FnCas12a-18nt  Repeat1 | F1: ATCGATTCCCTCACTCCTGCTCGGTGAA  R1: AGTAGACCAAGTCACTCTGGGGAACACGCC | TAATGCGC+CACAGGAT |
| B | T5 EXO-FnCas12a-18nt  Repeat2 | F2: CGATGTTCCCTCACTCCTGCTCGGTGAA  R2: TTATCTATAAGTCACTCTGGGGAACACGCC | TAATGCGC+CACAGGAT |
| B | T5 EXO-FnCas12a-18nt  Repeat3 | F3: TTAGGCTCCCTCACTCCTGCTCGGTGAA  R3: CATGACATAAGTCACTCTGGGGAACACGCC | TAATGCGC+CACAGGAT |
| C | FnCas12a-19nt  Repeat1 | F1: ATCGATTCCCTCACTCCTGCTCGGTGAA  R1: AGTAGACCAAGTCACTCTGGGGAACACGCC | TGGCACCT+AGCGAATG |
| C | FnCas12a-19nt  Repeat2 | F2: CGATGTTCCCTCACTCCTGCTCGGTGAA  R2: TTATCTATAAGTCACTCTGGGGAACACGCC | TGGCACCT+AGCGAATG |
| C | FnCas12a-19nt  Repeat3 | F3: TTAGGCTCCCTCACTCCTGCTCGGTGAA  R3: CATGACATAAGTCACTCTGGGGAACACGCC | TGGCACCT+AGCGAATG |
| D | T5 EXO-FnCas12a-19nt  Repeat1 | F1: ATCGATTCCCTCACTCCTGCTCGGTGAA  R1: AGTAGACCAAGTCACTCTGGGGAACACGCC | TGGCACCT+AGCGAATG |
| D | T5 EXO-FnCas12a-19nt  Repeat2 | F2: CGATGTTCCCTCACTCCTGCTCGGTGAA  R2: TTATCTATAAGTCACTCTGGGGAACACGCC | TGGCACCT+AGCGAATG |
| D | T5 EXO-FnCas12a-19nt  Repeat3 | F3: TTAGGCTCCCTCACTCCTGCTCGGTGAA  R3: CATGACATAAGTCACTCTGGGGAACACGCC | TGGCACCT+AGCGAATG |
| E | FnCas12a-20nt  Repeat1 | F1: ATCGATTCCCTCACTCCTGCTCGGTGAA  R1: AGTAGACCAAGTCACTCTGGGGAACACGCC | AAAGATAC+AGCGAATG |
| E | FnCas12a-20nt  Repeat2 | F2: CGATGTTCCCTCACTCCTGCTCGGTGAA  R2: TTATCTATAAGTCACTCTGGGGAACACGCC | AAAGATAC+AGCGAATG |
| E | FnCas12a-20nt  Repeat3 | F3: TTAGGCTCCCTCACTCCTGCTCGGTGAA  R3: CATGACATAAGTCACTCTGGGGAACACGCC | AAAGATAC+AGCGAATG |
| F | T5 EXO-FnCas12a-20nt  Repeat1 | F1: ATCGATTCCCTCACTCCTGCTCGGTGAA  R1: AGTAGACCAAGTCACTCTGGGGAACACGCC | TGGAGCTG+AGCGAATG |
| F | T5 EXO-FnCas12a-20nt  Repeat2 | F2: CGATGTTCCCTCACTCCTGCTCGGTGAA  R2: TTATCTATAAGTCACTCTGGGGAACACGCC | TGGAGCTG+AGCGAATG |
| F | T5 EXO-FnCas12a-20nt  Repeat3 | F3: TTAGGCTCCCTCACTCCTGCTCGGTGAA  R3: CATGACATAAGTCACTCTGGGGAACACGCC | TGGAGCTG+AGCGAATG |
| G | FnCas12a-21nt  Repeat1 | F1: ATCGATTCCCTCACTCCTGCTCGGTGAA  R1: AGTAGACCAAGTCACTCTGGGGAACACGCC | TAAGGCGA+AGCGAATG |
| G | FnCas12a-21nt  Repeat2 | F2: CGATGTTCCCTCACTCCTGCTCGGTGAA  R2: TTATCTATAAGTCACTCTGGGGAACACGCC | TAAGGCGA+AGCGAATG |
| G | FnCas12a-21nt  Repeat3 | F3: TTAGGCTCCCTCACTCCTGCTCGGTGAA  R3: CATGACATAAGTCACTCTGGGGAACACGCC | TAAGGCGA+AGCGAATG |
| H | T5 EXO-FnCas12a-21nt  Repeat1 | F1: ATCGATTCCCTCACTCCTGCTCGGTGAA  R1: AGTAGACCAAGTCACTCTGGGGAACACGCC | CGTACTAG+AGCGAATG |
| H | T5 EXO-FnCas12a-21nt  Repeat2 | F2: CGATGTTCCCTCACTCCTGCTCGGTGAA  R2: TTATCTATAAGTCACTCTGGGGAACACGCC | CGTACTAG+AGCGAATG |
| H | T5 EXO-FnCas12a-21nt  Repeat3 | F3: TTAGGCTCCCTCACTCCTGCTCGGTGAA  R3: CATGACATAAGTCACTCTGGGGAACACGCC | CGTACTAG+AGCGAATG |
| A | FnCas12a-22nt  Repeat1 | F4: TGACCATCCCTCACTCCTGCTCGGTGAA  R4: CGTTATGAAAGTCACTCTGGGGAACACGCC | CTGAAGCT+CACAGGAT |
| E | FnCas12a-22nt  Repeat2 | F4: TGACCATCCCTCACTCCTGCTCGGTGAA  R4: CGTTATGAAAGTCACTCTGGGGAACACGCC | AAAGATAC+AGCGAATG |
| A | FnCas12a-22nt  Repeat3 | F5: ACAGTGTCCCTCACTCCTGCTCGGTGAA  R5: TGCTCAACAAGTCACTCTGGGGAACACGCC | CTGAAGCT+CACAGGAT |
| B | T5 EXO-FnCas12a-22nt  Repeat1 | F4: TGACCATCCCTCACTCCTGCTCGGTGAA  R4: CGTTATGAAAGTCACTCTGGGGAACACGCC | TAATGCGC+CACAGGAT |
| F | T5 EXO-FnCas12a-22nt  Repeat2 | F4: TGACCATCCCTCACTCCTGCTCGGTGAA  R4: CGTTATGAAAGTCACTCTGGGGAACACGCC | TGGAGCTG+AGCGAATG |
| B | T5 EXO-FnCas12a-22nt  Repeat3 | F5: ACAGTGTCCCTCACTCCTGCTCGGTGAA  R5: TGCTCAACAAGTCACTCTGGGGAACACGCC | TAATGCGC+CACAGGAT |
| C | FnCas12a-23nt  Repeat1 | F4: TGACCATCCCTCACTCCTGCTCGGTGAA  R4: CGTTATGAAAGTCACTCTGGGGAACACGCC | TGGCACCT+AGCGAATG |
| G | FnCas12a-23nt  Repeat2 | F4: TGACCATCCCTCACTCCTGCTCGGTGAA  R4: CGTTATGAAAGTCACTCTGGGGAACACGCC | TAAGGCGA+AGCGAATG |
| C | FnCas12a-23nt  Repeat3 | F5: ACAGTGTCCCTCACTCCTGCTCGGTGAA  R5: TGCTCAACAAGTCACTCTGGGGAACACGCC | TGGCACCT+AGCGAATG |
| D | T5 EXO-FnCas12a-23nt  Repeat1 | F4: TGACCATCCCTCACTCCTGCTCGGTGAA  R4: CGTTATGAAAGTCACTCTGGGGAACACGCC | TAATGCGC+CACAGGAT |
| H | T5 EXO-FnCas12a-23nt  Repeat2 | F4: TGACCATCCCTCACTCCTGCTCGGTGAA  R4: CGTTATGAAAGTCACTCTGGGGAACACGCC | TGGAGCTG+AGCGAATG |
| D | T5 EXO-FnCas12a-23nt  Repeat3 | F5: ACAGTGTCCCTCACTCCTGCTCGGTGAA  R5: TGCTCAACAAGTCACTCTGGGGAACACGCC | TAATGCGC+CACAGGAT |
